# Sustained correction of hippocampal neurogenic and cognitive deficits after a brief treatment by Nutlin-3 in a mouse model of Fragile X Syndrome

**DOI:** 10.1101/2022.01.14.476402

**Authors:** Sahar Javadi, Yue Li, Jie Sheng, Lucy Zhao, Yao Fu, Daifeng Wang, Xinyu Zhao

## Abstract

**Background:** Fragile X syndrome (FXS), the most prevalent inherited intellectual disability and one of the most common monogenic form of autism, is caused by a loss of FMRP translational regulator 1 (FMR1). We have previously shown that FMR1 represses the levels and activities of ubiquitin ligase MDM2 in young adult FMR1-deficient mice and treatment by a MDM2 inhibitor Nutlin-3 rescues both hippocampal neurogenic and cognitive deficits in FMR1-deficient mice when analyzed shortly after the administration. However, it is unknown whether Nutlin-3 treatment can have long-lasting therapeutic effects.

**Methods:** We treated 2-month-old young adult FMR1-deficient mice with Nutlin-3 for 10 days and then assessed the persistent effect of Nutlin-3 on both cognitive functions and adult neurogenesis when mice were 6-month-old mature adults. To investigate the mechanisms underlying persistent effects of Nutlin-3, we analyzed proliferation and differentiation of neural stem cells isolated from these mice and assessed the transcriptome of the hippocampal tissues of treated mice.

**Results:** We found that transient treatment with Nutlin-3 of 2-month-old young adult FMR1-deficient mice prevents the emergence of neurogenic and cognitive deficits in mature adult FXS mice at 6-month of age. We further found that the long-lasting restoration of neurogenesis and cognitive function might not be mediated by changing intrinsic properties of adult neural stem cells. Transcriptomic analysis of the hippocampal tissue demonstrated that transient Nultin-3 treatment leads to significant expression changes in genes related to extracellular matrix, secreted factors, and cell membrane proteins in FMR1-deficient hippocampus.

**Conclusions:** Our data indicates that transient Nutlin-3 treatment in young adults leads to long-lasting neurogenic and behavioral changes through modulating adult neurogenic niche rather than intrinsic properties of adult neural stem cells. Our results demonstrate that cognitive impairments in FXS may be prevented by an early intervention through Nutlin-3 treatment.

## Introduction

Fragile X Syndrome (FXS) is the most common cause of inherited intellectual disability with prevalence rates estimated to be 1:5,000 in males and 1:8,000 in females [1]. FXS is one of the most common single-gene causes of autism spectrum disorder (ASD), with approximately 2 in 3 of male FXS patients being clinically diagnosed with ASD [2, 3]. FXS is mainly caused by an expansion of trinucleotide repeats (CGG) to over 200 repeats in the promoter region of the *FMR1* (FMRP translational regulator 1) gene which leads to transcriptional silencing of the gene with a subsequent reduction or absence of FMR1 (also known as FMRP, Fragile X Mental Retardation Protein) [4, 5]. FMR1 is a polyribosome-associated, brain-enriched, RNA-binding protein (RBP) that selectively targets specific mRNAs and regulates their translation, transport, and stability [4–8]. In addition, it has been shown that FMR1 is involved in histone modification and chromatin remodeling [9]. Hence, FMR1 is a multifunctional protein that could be involved in diverse biological processes.

FMR1 deficiency has been associated with numerous co-occurring conditions including, but not limited to, intellectual and emotional disabilities ranging from learning problems to mental retardation, and mood instability to autism [10]. Better understanding of the neurobiology and pathophysiology of FXS, together with advances in FXS animal models, has paved a way for the development of numerous targeted treatments for FXS. [11]. FMR1 is a multifunctional protein that regulates the expression of a large number of direct and indirect targets [8, 12, 13]. Despite the rich literature aiming to investigate short-term therapeutic outcomes of treatments, very few studies have evaluated potential long-lasting rescue effects of these treatments [11, 14]. Promising data from recent studies show that impairment of cognitive repertoire in FXS could be sustainably prevented by short-term pharmacological interventions [11, 15]. The long-lasting restoration of cognitive deficits of a rat model of FXS used in this study is associated with sustained rescue of both synaptic plasticity and altered protein synthesis These results have raised the question whether other reported interventions have the potential to exert long-lasting therapeutic effects.

Even though FMR1 is highly expressed in neurons, other cells have been implicated in FXS as well [16, 17]. Studies from our and other labs have shown that FMR1 regulates adult hippocampal neurogenesis [18–25]. Neurogenesis continuously occurs in at least two specific regions of the adult mammalian brain: the subventricular zone (SVZ) of the lateral ventricles and the subgranular zone (SGZ) of the dentate gyrus (DG) in the hippocampus [26]. Adult hippocampal neurogenesis is a multi-stage process, encompassing a number of developmental phases [26]. Activated neural stem cells (NSCs) or radial glia like cells (RGLs) generate intermediate neural progenitors (NPs) that subsequently differentiate into neuroblasts, immature neurons, and mature granule neurons (GCs) that finally integrate into existing circuits [27]. Adult hippocampal neurogenesis is implicated in many functional processes such as learning, memory, plasticity, and mood regulation [26] and are impaired in a number of neurological conditions including FXS [18–25]. Therefore, interventions aimed at regulating adult neurogenesis are being evaluated as potential therapeutic strategies in FXS[28]. We have previously shown that absence of FMR1 leads to increased NSC proliferation but reduced neuronal differentiation in young adult (2-month-old) *Fmr1* knockout (*Fmr1* KO) mice but reduced NSC proliferation and reduced neuronal differentiation in mature adult (6-month-old) *Fmr1* KO mice [19, 20]. At molecular levels, FMR1-deficient NSCs have elevated MDM2 (mouse double minute 2) protein levels and activities throughout adult ages. Treatment with Nutlin-3, a compound used for cancer clinical trial, specifically inhibits the interaction between MDM2 and its target proteins TP53 (Tumor Protein P53) and HDAC1 (Histone deacetylase 1) and rescues both adult hippocampal neurogenic and behavioral deficits in both young adult and mature adult FXS mice. However, it remains unknown whether Nutlin-3 can have long-lasting effects in FXS mice, which is a key question for therapeutic development.

In this study, we investigated whether Nutlin-3 treatment had sustained impact on neurogenesis and cognitive behaviors of FXS mice. We discovered that a transient treatment of young adult mice with Nutlin-3 led to long-lasting effect in both hippocampal neurogenesis and cognitive tasks in adult FXS mice. To our surprise, we found that the long-lasting effect of Nutlin-3 was not through modulating intrinsic properties of adult NSCs but rather through regulating the gene expression of the adult stem cell niche.

## Methods

### Study Design

The purpose of this study was to investigate the long-lasting effect of Nutlin-3 treatment on impaired neurogenesis and behavioral deficits in adult FXS mice. In addition, we aimed to determine the potential mechanisms underlying the sustained rescue effect by Nutlin-3. Based on our publications and power analysis, at least three biological replicates were used for each *in vitro* or *in vivo* biochemical and histological analysis, whereas a sample size of 9 to 21 per group was used for behavioral testing. The NPCs used for *in vitro* analyses were isolated from three pairs of *Fmr1* KO and WT littermates born to different parents and NPCs isolated from each animal was considered as a biological replicate. For drug treatment, animals were randomly assigned to treatment arms with approximately equivalent numbers in each group. All cell counting and behavioral analyses were performed by experimenters who were blind to the identity of the samples.

### Animal Studies

All animal procedures were performed according to protocols approved by the University of Wisconsin-Madison Care and Use Committee. All mice were on C57B/L6 genetic background. The crossing and genotyping of these mice were carried out as described previously [19, 20]. Briefly, the *Fmr1* KO;*Nestin*-GFP mice (*Fmr1^−y^;Nestin*-GFP) were created by crossing female *Fmr1* heterozygous KO mice (*Fmr1*^+/−^) [29] with homozygous *Nestin*-GFP transgenic males [30]. Generation of FMR1 inducible conditional mutant mice (*Fmr1^loxP/y^;Nestin-CreER^T2^;Rosa26-tdT* or *cKO;Cre;Ai14*) and tamoxifen injections to induce recombination were performed as described [21]. To induce recombination, mice (6-week old) received tamoxifen (160 mg/kg; Sigma-Aldrich) daily for 5 days as described [21]. Nutlin-3 (10 mg/kg) was dissolved in dimethyl sulfoxide, given to 2-month-old mice through intraperitoneal injections every other day for five injections, and sacrificed at either 4-months after the last injection. For in vivo differentiation analysis, mice also received four BrdU injections (100 mg/kg) within 12 hours at one month before sacrifice.

### Tissue Preparation and Immunohistochemistry

Brain tissue processing and histological analysis of mouse brains were performed as described in our publications[18–21, 31]. Briefly, mice were euthanized by intraperitoneal injection of a mixture of ketamine/xylazine/acepromazine followed by transcardiac perfusion with saline and then 4% paraformaldehyde (PFA). Brains were dissected out, post-fixed overnight in 4% PFA, and equilibrated in 30% sucrose. Brain sections of 40 μm thickness were generated using a sliding microtome and stored in a −20 °C freezer as floating sections in cryoprotectant solution (glycerol, ethylene glycol, and 0.1Mphosphate buffer (pH 7.4), 1:1:2 by volume). We performed immunohistological analysis on 1-in-6 serial floating brain sections (240 μm apart). After staining with primary and fluorescent secondary antibodies, sections were counterstained with DAPI (1:1000; Roche Applied Science) and then mounted, coverslipped, and maintained at 4°C in the dark until analysis.

#### The primary antibodies used were

Chicken-anti-GFP (1:1000, Invitrogen, A10262), rabbit anti-MCM2 (1:500, Cell Signaling, 4007), rabbit anti-GFAP (1:1000, Dako, Z0334), mouse anti-GFAP (1:1000, Millipore, MAB360), mouse anti-NeuN (1:500, Millipore, MAB377), rat anti-BrdU (1:1000, Abcam, ab6326)

#### Fluorescent secondary antibodies used were

goat anti-mouse 568 (1:1000, Invitrogen, A11004), goat anti-rat 568 (1:1000, Invitrogen, A11077), goat anti-rabbit 568 (1:1000, Invitrogen, A11011), goat anti-rabbit 647 (1:1000, Invitrogen, A21245), goat anti-mouse 647 (1:1000, Invitrogen, A21235), goat anti-chicken 488 (1:1000, Invitrogen, A11039), goat anti-mouse 488 (1:1000, Invitrogen, A11029).

### *In Vivo* Cell Quantification

Quantitative analyses of adult neurogenesis were carried out using unbiased stereology method through the use of a Stereo Investigator software (MBF Biosciences) as described [19, 20, 32]. Briefly, Z-stack images (2μm interval) were acquired using an AxioImager Z2 ApoTome confocal microscope (Plan-APOCHOROMAT, 20X, numerical aperture=0.8; Zeiss). The measured thickness of the sections were ~30 μm. The cell numbers were quantified by random sampling one in six coronal serial sections (240 μm apart) encompassing the entire hippocampus, with 3 μm guard zones on each side. Schaeffer’s coefficient of error (CE) <0.1 was required for each type of cell quantification. The experimenter was blinded to the identity of the samples. The total numbers of GFP+GFAP+ cells in the dentate gyrus of each animal were counted. Then, the percentage of activated and proliferating NSCs were determined by co-localizing of GFP+GFAP cells with cell cycle marker, MCM2. Cell lineage analysis was performed as described [19]. Briefly, at least 100 BrdU^+^ cells in the Z-stacks of each mouse were randomly selected, and their co-localization with cell-lineage marker, NeuN, was determined using Stereo Investigator.

### Novel Location Recognition Test

This test measures spatial memory through an evaluation of the ability of mice to recognize the new location of a familiar object with respect to spatial cues. The experimental procedure was carried out as described previously [19, 20]. Briefly, mice were handled for approximately 5 min a day for a maximum of 5 days prior to the experiment. Testing consisted of five 6-min trials, with a 3-min intertrial interval between each trial. All procedures were conducted during the light cycle of the animal between 10 a.m. and 4 p.m. Before the trial session, mice were brought into the testing room and were allowed to acclimate for at least 30 minutes. During the intertrial interval, the mouse was placed in a holding cage, which remained inside the testing room. In the first trial (Pre-Exposure), each mouse was placed individually into the center of the otherwise empty open arena (38.5cm Long×38.5cm wide, and 25.5cm high walls) for 6 min. For the next three trials (Sample Trials 1-3), two identical objects were placed equidistantly from the arena wall in the corners against the wall with the colored decal. Tape objects to the floor of the arena. Then, each mouse was placed individually into the center of the arena and was allowed to explore for 6 minutes. At the end of the trial, the mouse was removed and returned to the home cages for 3 minutes. In the last trial (Test), one of the objects was moved to a novel location, and the mouse was allowed to explore the objects for 6 minutes, and the total time spent exploring each object was measured. During the test phase, exploration time was defined as any investigative behavior (i.e., head orientation, climbing on, sniffing occurring within < 1.0 cm) or other motivated direct contact occurring with each object. To control for possible odor cues, objects were cleaned with 70% ethanol solution at the end of each trial and the floor of the arena wiped down to eliminate possible scent/trail markers. During the test phase, two objects were wiped down prior to testing so that the objects would all have the same odor. Based on a previous study [33], the discrimination index was calculated as the percentage of time spent investigating the object in the new location minus the percentage of time spent investigating the object in the old location: discrimination index = (Novel Location exploration time/total exploration time × 100) – (old Location exploration time/total exploration time × 100). A higher discrimination index is considered to reflect greater memory retention for the novel location object. All experiments were videotaped and scored by scientists who were blinded to experimental conditions to ensure accuracy.

### Novel Object Recognition Test

This test is based on the natural propensity of rodents to preferentially explore novel objects over familiar ones. The experimental procedure was carried out as described previously [19, 20]. Briefly, mice were handled for approximately 5 min a day for a maximum of 5 days prior to the experiment. The test was conducted during the light cycle of the animal between 10 a.m. and 4 p.m. Before the trial or test phase, mice were brought into testing room and were allowed to acclimate for at least 30 minutes. On the first day, mice were habituated for 10 min to the V-maze, made out of black Plexiglas with two corridors (30cm Long × 4.5cm wide, and 15cm high walls) set at a 90-degree angle, in which the task was performed. On the second day, mice were put back in the maze for 10 min, and two identical objects were presented. 24 hours later, one of the familiar objects was replaced with a novel object, and the mice were again placed in the maze and were allowed to explore for 10 min, and the total time spent exploring each of the two objects (novel and familiar) was measured. During the test phase, the novel and familiar objects were wiped down prior to testing so that the objects would all have the same odor and exploration time was defined as the orientation of the nose to the object at a distance of less than 2 cm. The discrimination index was calculated as the difference between the percentages of time spent investigating the novel object and the time spent investigating the familiar objects: discrimination index = (novel object exploration time/total exploration time × 100) – (familiar object exploration time/total exploration time × 100). A higher discrimination index is considered to reflect greater memory retention for the novel object. All experiments were videotaped and scored by scientists who were blinded to experimental conditions to ensure accuracy.

### Adult NSPC Isolation and Analyses

NSPCs were isolated from the pooled DG tissue dissected from two 6-month-old male mice using our published method [34] [20]. NSPCs were cultured as described previously [34]. Proliferation and differentiation of NPCs were analyzed as described [19]. We used only early passage cells (between passages 4 and 10) and only the same passage numbers of wild-type and *Fmr1* KO cells. For each experiment, triplicate wells of cells were analyzed, and results were averaged as one data point (n = 1). At least three independent biological replicates were used (n = 3) for statistical analyses.

The primary antibodies used were Mouse-anti-Tuj1 (1:1000, Covance, 435P) and rat anti-BrdU (1:3000, Abcam, ab6326)

Fluorescent secondary antibodies used were goat anti-mouse 568 (1:2000, Invitrogen, A11004) and goat anti-rat 568 (1:2000, Invitrogen, A11077)

### RNA Isolation and RNA-seq

Freshly dissected hippocampal tissue was immediately frozen on dry ice. For RNA isolation, Trizol was added to frozen tissue followed by homogenization using a Polytron (vendor?). RNA was isolated from TRIzol samples using the TRIzol Reagent following the manufacturer’s instructions. RNA quality assessment, library construction, library quality control and sequencing were performed by Novogene Bioinformatics Institute (Sacramento, CA, USA). Briefly, the quality, size, and concentration of the isolated RNA were analyzed using agarose gel electrophoresis, nanodrop, and an Agilent 2100 Bioanalyzer. Twelve cDNA libraries were constructed with three biological replicates for each condition: WT-veh, KO-veh, WT-Nut3, and KO-Nut3. Messenger RNA was purified from total RNA using poly-T oligo-attached magnetic beads. After fragmentation, the first strand cDNA was synthesized using random hexamer primers followed by the second strand cDNA synthesis. The library was ready after end repair, A-tailing, adapter ligation, size selection, amplification, and purification (Novogene, Sacramento, CA, USA). Library quality was assessed by Qubit 2.0, Agilent 2100, and qPCR. The libraries were clustered and sequenced on an Illumina Novaseq 6000 on an S4 flow cell. The 150 bp paired-end reads were generated after clustering of the index-coded samples. About 20-30 million reads were obtained for each sample.

### Bioinformatics Analysis

FastQC was used to perform quality check of .fastq reads. Paired end reads were mapped to reference genome (mm10) using STAR. Raw count matrix was normalized by correcting for library size using DESeq2 R package (**Additional file 1**). Differential expression analysis was performed by using Dseq2 R package (**Additional file 1**). Adjusted P-value < 0.05 were used as cutoffs for differential expression. GO term enrichment was analyzed using Enricher (https://maayanlab.cloud/Enrichr)[35] and plotted using GOplot[36]. Transcription factor enrichment analysis and network were performed using ChEA3 (https://maayanlab.cloud/chea3)[37] and average integrated ranks across all libraries. Submission of RNA-seq data to Gene Expression Omnibus (GEO) is in process. $

### Real-time PCR Assay

Real-time PCR was performed using standard methods as described [19]. The first-strand cDNA was generated by reverse transcription with Oligo (dT) primer (Roche). To quantify the mRNA levels using real-time PCR, aliquots of first-stranded cDNA were amplified with gene-specific primers and Power SYBR Green PCR Master Mix (Bio-Rad) using a Step-1 Real-Time PCR System (Applied Biosystems). The PCR reactions contained 1μg of cDNA, Universal Master Mix (Applied Biosystems), and 10μM of forward and reverse primers in a final reaction volume of 20μL. The data analysis software built in with the 7300 Real-Time PCR System calculated the mRNA level of different samples. The sequences of primers used for Real-time PCR reactions in mouse species are listed in **Additional file 2: Table S1**.

### Western Blotting Analyses

Protein samples were separated on SDS-PAGE gels (Bio-Rad), transferred to PVDF membranes (Millipore), and incubated with primary antibodies. The antibodies include p-MDM2(Ser166,1:1000, Novus Biologicals, NBP1-51396), HDAC1(1:1000, BioVersion, 3601-30), Acetyl-H3 (1:1000, Millipore, 06-599), Histone H3 (1:1000, Cell Signaling, 9715S), and GAPDH (1:5000, Thermo Scientific, MA5-15738). After incubation with fluorescence-labeled secondary antibodies (Li-CoR), the membranes were imaged using Li-CoR and quantification was performed using Image Studio Lite software. The amount of loading protein (20μg) was determined by the linear range of the target proteins (10μg-40μg) using Li-CoR system as previous described [19]. At least three independent blots were used for statistical analysis.

### Statistical Analysis

All experiments were randomized and blinded to scientists who performed quantification. Statistical analysis was performed using ANOVA and Student’s t-test, unless specified, with the GraphPad Prism software 9. Two tailed and unpaired t-test was used to compare two conditions. Two-way ANOVA with Tukey’s post hoc analysis was used for analyzing multiple groups. All data were shown as mean with standard error of mean (mean ± SEM). Probabilities of P<0.05 were considered as significant.

## RESULTS

### Transient Nutlin-3 treatment has long-lasting rescue effect on impaired hippocampal neurogenesis in *Fmr1* KO mice

We have previously shown that young adult (2-month-old) *Fmr1 KO* mice exhibited elevated NSC activation and impaired neurogenesis and mature adult (6-month-old) *Fmr1* KO mice exhibited reduced NSC activation and impaired neurogenesis in the hippocampus, which can be normalized to WT levels immediately after a 10-day treatment by a specific MDM2 inhibitor Nutlin-3 [19, 20]. To assess the potential of MDM2 inhibition as a therapeutic treatment, a critical question remained is whether Nutlin-3 treatment has long-lasting rescue effects. We thus decided to investigate whether a transient Nutlin-3 treatment could have persistent therapeutic effect on NSC activation and adult neurogenesis in FXS mouse models.

We crossed *Fmr1* mutant mice with *Nestin*-GFP (green fluorescent protein) mice in which GFP expression is driven by the promotor of a neural stem and progenitor cell marker NESTIN to create the *Fmr1* KO;*Nestin-GFP* double transgenic mice as described previously [19]. We treated 2-month-old *Fmr1 KO* (*Fmr1^−/y^;Nestin-GFP*) and littermate wild-type (*Fmr1*^+/y^;*Nestin-GFP*) mice with either vehicle or Nutlin-3 (10 mg/kg) every other day over 10 days (total 5 injections) as we have done previously [19, 20] and analyzed them at 4-months after the last injections, when the mice were 6-month-old (**Fig. 1a**). Glial fibrillary acidic protein (GFAP) is a radial glia marker expressed in both quiescent and activated adult hippocampal NSCs (**Fig. 1b**) [38]. To determine activation of NSCs, we used the cell cycle marker minichromosome maintenance complex component 2 (MCM2; **Fig. 1b**). We quantified the percentage of activated (GFP^+^GFAP^+^MCM2^+^) NSCs over total (GFP^+^GFAP^+^) NSCs in *Fmr1* KO and WT mice. We found that *Fmr1* KO mice treated with vehicle at 2-month of age exhibited reduced NSC activation at 6-month of age compared to WT with the same vehicle treatment (**Fig. 1c**), which is consistent with our previous finding on 6-month-old *Fmr1* KO mice [20]. Similarly to what we have published before [19, 20], Nutlin-3 treatment had no significant effect on WT mice (**Fig. 1c**). In contrast, *Fmr1* KO mice treated with Nutlin-3 at 2-month of age showed no significant difference in NSCs activation at 6-month of age compared to WT mice treated with either vehicle or Nutlin-3 (**Fig. 1c**). Therefore, a 10-day transient Nutlin-3 treatment of young adult *Fmr1* KO mice has long-lasting rescue effect on adult hippocampal NSC activation, which persists for at least 4 months.

**Fig. 1:**
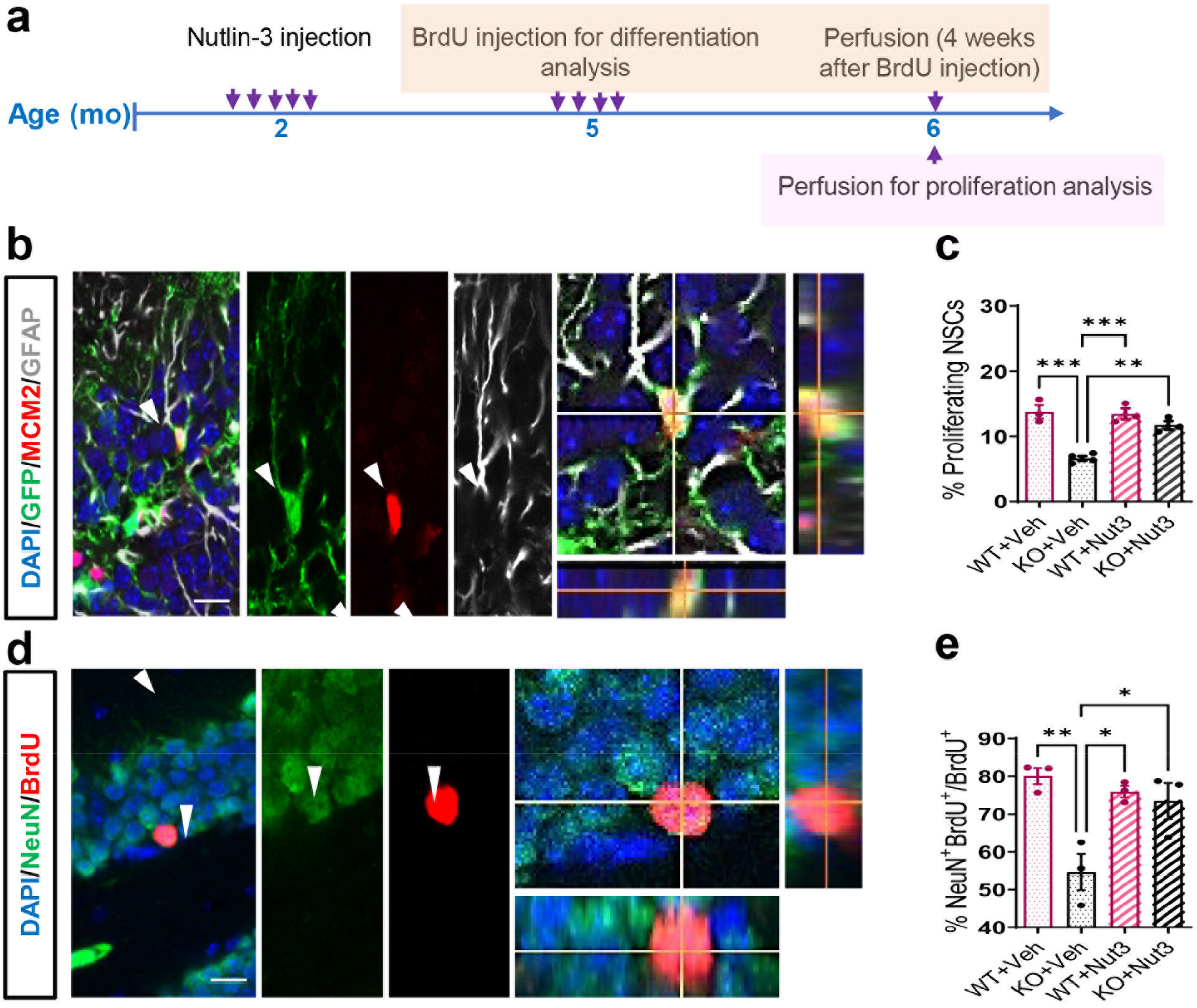
Transient treatment with Nutlin-3 has long-lasting rescue effect on impaired hippocampal neurogenesis in FMR1-deficient mice. **a** Experimental scheme for assessing hippocampal neurogenesis in *Fmr1* KO and WT mice treated with Nutlin-3 or vehicle. **B** Sample confocal images used for identifying NSCs (GFP+GFAP+) and proliferating NSPCs (GFP+GFAP+MCM2+) in the dentate gyrus of adult *Fmr1* KO and WT mice bred into a *Nestin*-GFP mouse background. Scale bar, 20 μm. **c** Comparison of the percentage of activated NSPCs among total NSPCs in the DG of *Fmr1* KO and WT mice with or without Nutlin-3 treatment (n = 3 or 4 per group). **d** Sample confocal images to identify new mature neurons (NeuN+BrdU+) in the dentate gyrus of *Fmr1 KO* and WT mice. Scale bar, 20 μm. **e** Comparison of the percentage of mature neurons among BrdU+ cells in DG of *Fmr1* KO and WT mice with or without Nutlin-3 treatment (n = 3 per group). *P < 0.05; **P < 0.01 ***P < 0.001. Data are presented as means ± SEM.

We then assessed whether the therapeutic effect of Nutlin-3 on neuronal differentiation [19, 20] could persist long after treatment. Thus, we injected 2-month-old *Fmr1* KO (*Fmr1^−/y^*) mice and WT (*Fmr1^+/y^*) littermates with Nutlin-3 as described above (**Fig. 1a**) [19]. At 5-month of age, the mice received four injections of a synthetic thymidine analogue bromodeoxyuridine (BrdU) over a 12-hour period to pulse label proliferating NSCs and progenitors in the adult DG and were sacrificed at 4 weeks after BrdU injections (6-month of age) for differentiation analysis (**Fig. 1a**) [19]. To identify the fate of the BrdU-labeled NSPCs, we performed co-immunostaining using antibodies against mature neuronal marker NeuN (neuronal nuclei antigen or RBFOX3) and BrdU and quantified the percentage of neuronal differentiation (BrdU^+^NeuN^+^/BrdU^+^) (**Fig. 1d**). We found that *Fmr1* KO mice treated with vehicle showed a significant reduction in neuronal differentiation compared to WT counterparts treated with either vehicle or Nutlin-3 (**Fig. 1e**), which is consistent as what has been reported previously [20]. In contrast, 6-month-old *Fmr1* KO mice treated with Nutlin-3 at 2-month of age showed elevated neuronal differentiation level comparable to that of the WT mice (**Fig. 1e**). Nutlin-3 administration did not show significant effects on neurogenesis of WT mice (**Fig. 1e**). Therefore, transient treatment of *Fmr1* KO mice with Nutlin-3 at young adult ages could prevent impairment of neuronal differentiation at mature adult mice. In summary, our findings have revealed a long-lasting therapeutic effect of Nutlin-3 on impaired NSC activation and neuronal differentiation in a FXS mouse model.

### Transient Nutlin-3 treatment has endured corrective effect on cognitive deficit in FXS mouse models

We have previously shown that *Fmr1* KO mice exhibited deficits in hippocampus-dependent cognitive functions [19–21, 39] and Nutlin-3 treatment reversed impaired spatial learning assessed by a novel location recognition test (NLR) and defective cognitive function assessed by a novel object recognition (NOR) test at one month after treatment [19, 20]. Since our current study has revealed a persistent therapeutic effect of Nutlin-3 on impaired NSCs activation and neurogenesis in *Fmr1* KO mice (**Fig. 1**), we decided to investigate whether the Nutlin-3-dependent restoration of cognitive deficit is also long-lasting.

First, to confirm that selective deletion of FMR1 from NSCs in young adult mice leads to long-lasting impaired performances on hippocampus-dependent learning tasks, we generated tamoxifen-inducible *Fmr1* conditional knockout (cKO;CreER^T2^;Ai14) triple transgenic mice by crossing *Fmr1*-floxed (*Fmr1^f/f^*, or cKO) mice with inducible *Nestin* promotor-driven Cre transgenic mice (*Nes-CreER^T2^*) and *Rosa26-STOP-tdTomato* (Ai14) reporter mice as described previously [19] (**Additional file 3: Fig. S1a**). We found that targeted deletion of FMR1 from NSCs and their progenies at 2-month of age led to learning deficits at 6-month of age (**Additional file 3: Fig. S1b-d**), which corroborated our previous findings [19, 21]. More importantly, transient treatment of *cKO;CreER^T2^;Ai14* mice with Nutlin-3 at 2-month of age led to restoration of cognitive function when assessed at 6-month of age (**Additional file 3: Fig. S1b-d**). Therefore transient Nutlin-3 treatment in young adult mice with selective deletion of FMR1 from adult new neurons has long lasting effect.

We then treated 2-month-old *Fmr1* KO mice and their WT littermates with either vehicle or Nutlin-3 and analyzed their behaviors 4 months later (**Fig. 2a**). Consistent with our previous findings, *Fmr1* KO mice treated with vehicle exhibited impaired performance in spatial learning on the NLR test and defective learning on the NOR test (**Fig. b-e**)[20]. In contrast, Nutlin-3 administration rescued the impaired performances of *Fmr1* KO mice in both NLR (**Fig. 2c**) and NOR (**Fig. 2e**) to the wild-type levels without significant effect on wild-type mice (**Fig. 2c,e**). Therefore, a transient Nutlin-3 treatment of FMR1-deficient mice at young adulthood could rescue impaired cognitive performance for at least 4 months.

**Fig. 2:**
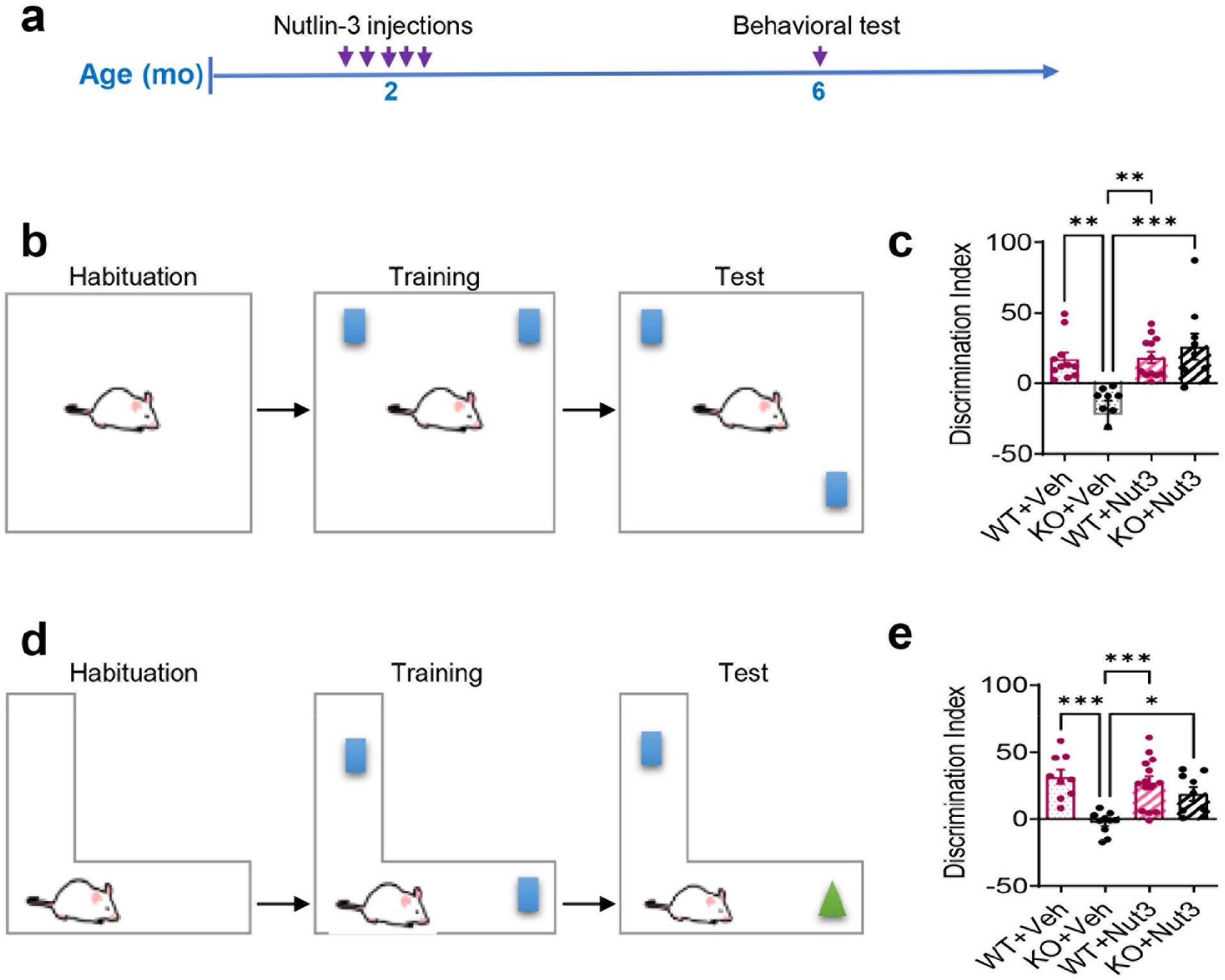
Transient treatment with Nutlin-3 has long-lasting rescue effect on cognitive deficits in FMR1-deficient mice. **a** Experimental scheme for analyzing cognitive performances in *Fmr1* KO and WT mice treated with Nutlin-3 or vehicle. **b** Schematic of novel location test for assessing spatial learning. **c** Beneficial effects of Nutlin-3 treatment on spatial memory deficits in *Fmr1* KO mice sustained at least 4-months after injection (n = 8 to 13mice per group). **d** Schematic of the novel object recognition test. **e** Therapeutic effects of Nutlin-3 treatment on deficits in the novel object recognition test in *Fmr1* KO mice last at least for 4-months after treatment cessation (n = 8 to 13 mice per group). *P < 0.05; **P < 0.01 ***P < 0.001. Data are presented as means ± SEM.

### Transient treatment with Nutlin-3 does not have persistent effect on intrinsic properties of adult neural stem/progenitor cells

We next sought to reveal the molecular mechanisms that are associated with Nutlin-3-induced enduring rescue of impaired hippocampal neurogenesis and related cognitive functions. Adult NSCs and adult hippocampal neurogenesis are regulated by both intrinsic and extrinsic factors [40]. To determine whether Nutlin-3 treatment acted through NSC intrinsic pathways, we decided to analyze the neural stem/progenitor cells (NSPCs) isolated from the hippocampus of 6-month-old *Fmr1* KO and littermate WT mice treated with either vehicle or Nutlin-3 at 2-month of age (**Fig. 3a**) using our published methods [34]. We used BrdU pulse labeling to assess NSPC proliferation and found that *Fmr1 KO* NSPCs exhibited reduced BrdU incorporation rate compared to WT NSPCs **(Fig. 3b,d)**, consistent with our published results on NSPCs isolated from 6-month-old *Fmr1 KO* mice [20]. Surprisingly, Nutlin-3 treatment did not rescue the impaired proliferation of *Fmr1* KO cells **(Fig. 3d)**. We then assessed NPSC neuronal differentiation using an antibody for immature neuron, βIII-tubulin (Tuj1) and found that NSPCs isolated from *Fmr1* KO mice at 4-months after either Nutlin-3 or vehicle injection exhibited similarly reduced neuronal differentiation (**Fig. 3c,e**). We have previously shown that NSCs isolated from 6-month old mice exhibited elevated phosphorylated MDM2 (P-MDM2, the active form of MDM2), acetylated Histone H3, and HDAC1 levels [20] We therefore assessed the levels of these proteins and found that NSPCs isolated from KO mice exhibited increased P-MDM2 protein levels, elevated H3 acetylation, and reduced HDAC1 levels, consistent to our published results (**Fig. 3f-k**) [20]. More importantly, treatment of Nutlin-3 at 2-month of age did not alter the levels of these proteins in the NSPCs isolated from 6-month-old mice (**Fig. 3f-k**). Therefore, these data revealed that long-lasting effect of Nutlin-3 on hippocampal NSPCs was unlikely due to intrinsic changes in NSPCs.

**Fig. 3:**
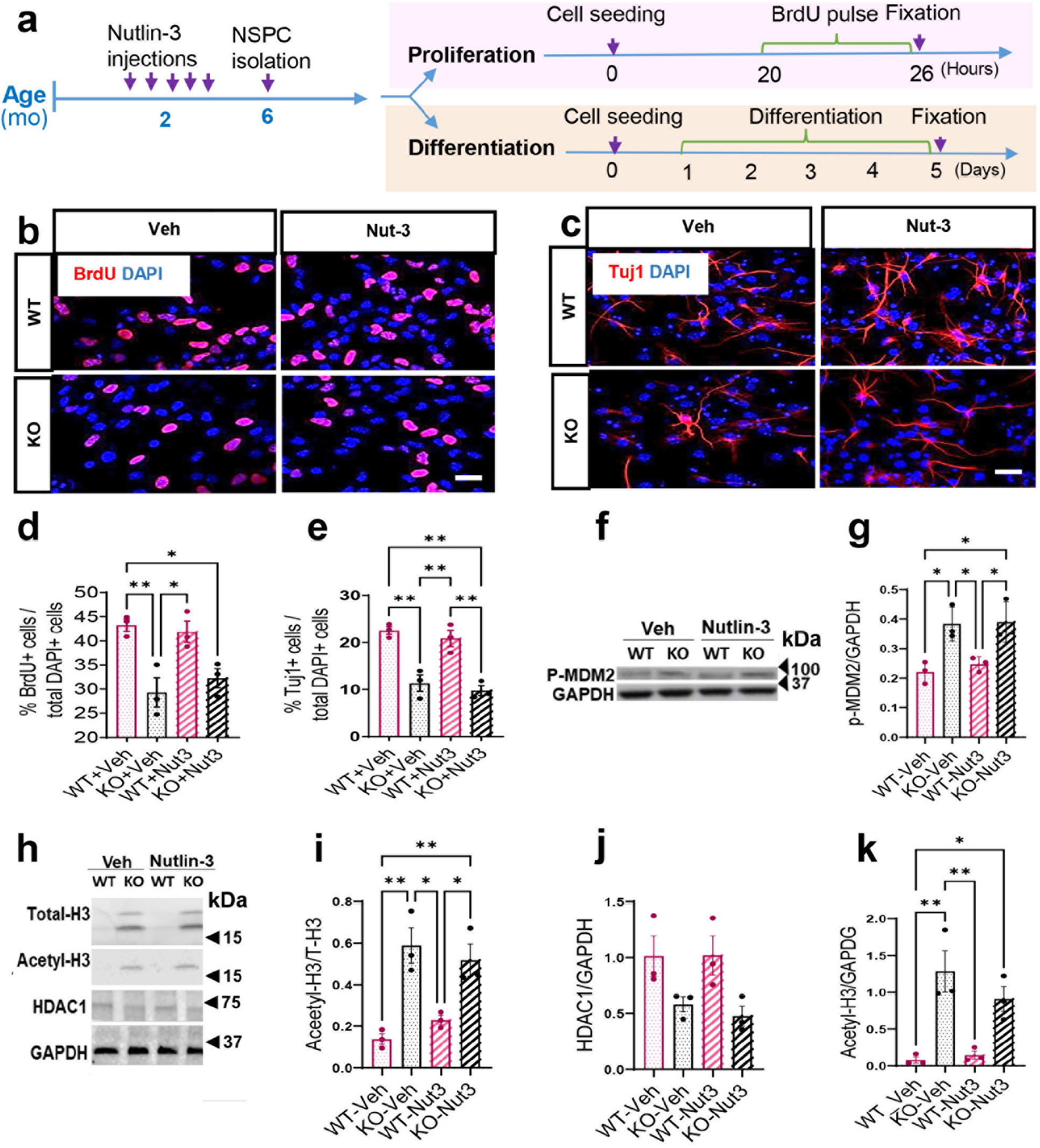
Transient treatment with Nutlin-3 does not have persistent effect on intrinsic properties of adult neural progenitor cells. **a** Experimental scheme for analyzing proliferation and differentiation of hippocampal NSPCs isolated from *Fmr1* KO and WT mice treated with Nutlin-3 or vehicle. **b** Sample images of proliferating NSPCs pulse labeled with thymidine analog, BrdU followed by immunohistology for in vitro quantification assay. Red, BrdU; blue, DAPI; scale bar, 20 μm. **c** Sample images of differentiating NSPCs assessed by immunohistological detection of a neuronal marker Tuj1^+^ for in vitro quantification of NSPC neuronal differentiation. Red, Tuj1; blue, DAPI; scale bar, 20 μm. (**d, e**) Nutlin-3 treatment did not rescue impaired proliferation (**d**) and neuronal differentiation (**e**) of hippocampal NSPCs isolated from *Fmr1* KO and WT mice 4-months after injection (n = 3). (**f, g**) Western blot analysis of P-MDM2 levels in isolated NSPCs isolated from *Fmr1* KO and WT 4-months after Nutlin-3 or vehicle treatment (n = 3). GAPDH was used as loading control. (**h, k**) Western blot analysis of total histone H3, acetylated histone H3, and HDAC1 levels in NSPCs isolated from *Fmr1* KO and WT mice 4-months after Nutlin-3 or vehicle treatment (n = 3). Glyceraldehyde-3-phosphate dehydrogenase (GAPDH) was used as loading control. *P < 0.05; **P < 0.01 ***P < 0.001. Data are presented as means ± SEM.

### Transient Nutlin-3 treatment leads to significant and specific gene expression changes in *Fmr1* KO hippocampus

Since transient Nutlin-3 treatment did not have significant rescue effect on NSPCs isolated 4-months later, we reckoned that Nutlin-3 might exert its long-lasting impact on hippocampal neurogenesis through modulating stem cell niche in the hippocampus. To explore the potential regulatory mechanisms, we injected 2-month-old *Fmr1* KO and WT control mice with Nutlin-3 or vehicle and harvested the hippocampi at 4-months post-treatment for transcriptomic analysis in triplicates (**Fig. 4a**). Total sequencing reads generated for each sample were between 21 and 26 million (21×10^6^ <TRs< 26×10^6^) (**Additional file 2: Table S2**). More than 94% reads were uniquely mapped to the mouse genome, which corresponds to more than 25×10^3^ genes (**Additional file 2: Table S2,3**). We evaluated the distribution of read counts across the samples and found that the overall density distribution of raw log-intensities exhibited a highly consistent pattern (**Additional file 3: Fig. S2**).

**Fig. 4:**
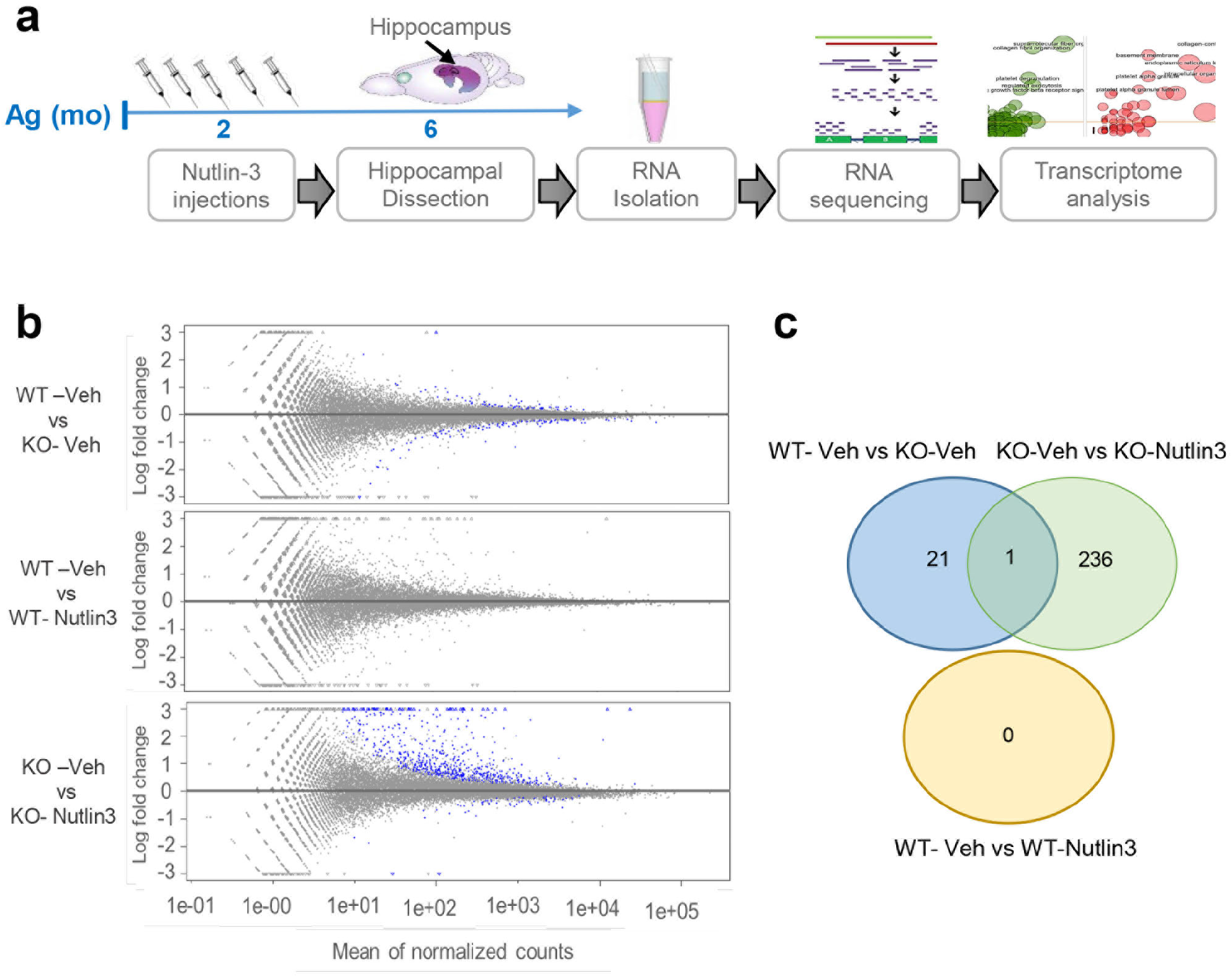
Transient treatment with Nutlin-3 leads to long-lasting gene expression changes in the hippocampus of FMR-deficient mice. **a** Experimental time line for sample collection and transcriptomic profiling of the hippocampal tissue of *Fmr1* KO and WT mice injected with Nutlin-3 or vehicle (n = 3 per group). **b** M-A plot of M (log ratio) and A (mean average) displaying log2 fold-change of genes compared with mean expression levels of all genes with log2 fold-change thresholds between −3 and 3. The genes identified differential expression (adjusted P < 0.05) are indicated as blue dots. **c** Venn diagram showing overlap patterns of differentially expressed genes between different experimental groups.

Next, we performed differential expression analysis to identify significantly deregulated genes among the four experimental groups. Adjusted P-value < 0.05 were used as cutoffs for differential expression. We identified no differentially expressed genes (DEGs) between vehicle-treated and Nutlin-3-treated WT mice, consistent with a lack of effect by Nutlin-3 on WT mice shown in our published results [19, 20] and current neurogenic and behavioral data (**Fig. 1–3, Fig. 4b,c; Additional file 2: Table S4**). We identified 21 DEGs between vehicle-treated *Fmr1 KO* (KO-Veh) and WT mice (WT-Veh), of which 13 DEGs were downregulated and 8 genes were upregulated (**Fig. 4c; Additional file 2: Table S4**). Surprisingly, we found that the most significant gene expression changes were between vehicle treated and Nulin-3-treated *Fmr1* KO mice (KO-Veh vs KO-Nut3) (**Fig. 4c; Additional file 2: Table S4**). Out of a total 237 DEGs (KO-Veh vs KO-Nut3), 6 gene were downregulated and 231 genes were upregulated in Nutlin-3 treated KO mice (**Fig. 4c; Additional file 2: Table S4**). Only 1 DEG, gene Gm21887 or *Erdr1* (erythroid differentiation regulator 1), was shared between these any two groups of DEGs (**Fig. 4c; Additional file 2: Table S4**). Therefore transient Nutlin-3-triggered significant gene expression changes in KO mice but did not make gene expression in KO mice to be more like that in WT mice.

To understand the biological significance of DEGs found in *Fmr1* KO mice treated with Nutlin-3, we performed Gene Ontology (GO) analysis using three categories of term analysis: Biological Pathway, Cell Component and Molecular Function (**Additional file 2: Table S5**). We generated circle plots to demonstrate specific enrichment and the directionality of the gene expression changes within each GO category (**Fig. 5a**). The DEGs were generally enriched for extra cellular matrix, cell membrane proteins and secreted factors (**Fig. 5a; Additional file 2: Table S5**) known to be key components of adult neurogenic niche [41–44], including the well-known BMP and TGFß signaling pathway [45] and IGF2 pathway [46]. In each GO category there was a robust upregulation of the DEGs for the enriched terms (**Fig. 5a**).

**Fig. 5:**
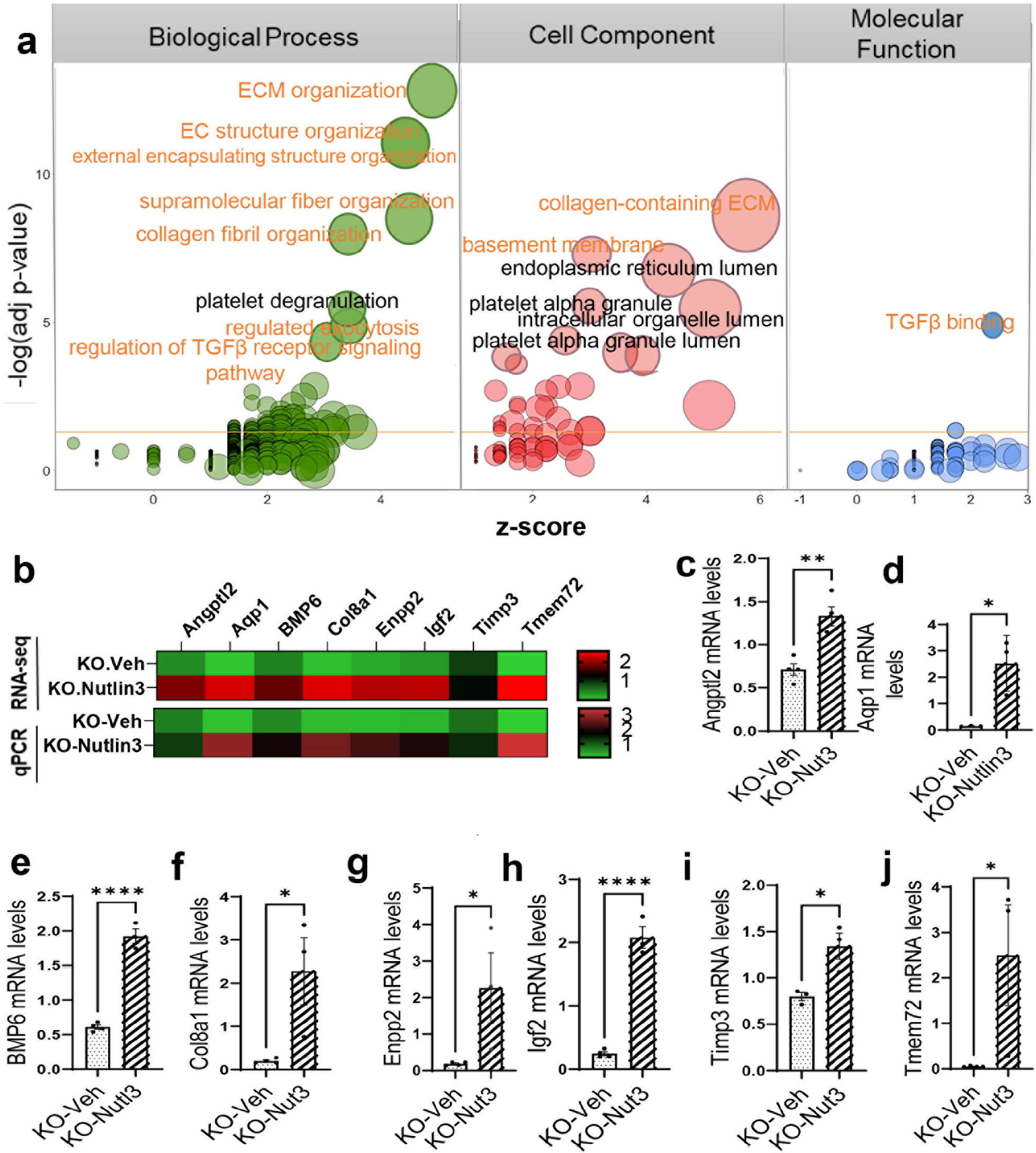
DEGs in FMR-deficient mice treated with Nutlin-3 were enriched for gene associated adult neural stem cell niche regulation. **a** Bubble plots for gene ontology (GO) analysis showing enriched terms identified with Enricher for DEGs between *Fmr1* KO treated with Nutlin-3 or Vehicle. Three different categories of GO analysis results are shown. The size of bubbles indicates the number of genes. The *x* axis indicates *z* score (negative = downregulated in Nutlin-3 treated *Fmr1* KO mice; positive = upregulated in Nutlin-3 treated *Fmr1* KO mice). The *y* axis indicates negative logarithm of adjusted *P* value from GO analysis (higher = more significant). ECM organization, membrane proteins and secreted factors are top hits in each GO category. **b** Heat map of transcriptional changes of selected DE genes between Nutlin-3 and vehicle-treated *Fmr1* KO mice, revealed by DESeq2 (n = 3) and qPCR analysis (n= 4). Red and green represent upregulation and downregulation, respectively. (**c-j**) Quantitative PCR analysis to validate a subset of DEGs in each GO category including Angptl2 (c), Aqp1 (d), Bmp6 (e), Col8a1 (f), Enpp2 (g), Igf2 (h), Tmp3 (i), and Tmem72 (j) (n = 3/condition). The mRNA levels of Glyceraldehyde-3-phosphate dehydrogenase (GAPDH) was used as the internal control. *P < 0.05; **P < 0.01 ***P < 0.001. Data are presented as means ± SEM.

Because the enriched terms have shown strong potential in stem cell regulation through modulating stem cell niche [41–44], we next selected a number of candidate DEGs from each group and validated their differential expression in KO-Nut3 compared to KO-Veh using quantitative polymerase chain reaction (qPCR) analysis. Administration of Nutlin-3 in *Fmr1* KO mice led to significant changes in the expression levels of genes associated to extracellular matrix (*Col8a1* and *Timp3*), cell membrane (*Aqp1* and *Tmem72*) and secreted factors (*Angptl2, Enpp2, BMP6* and *Igf2*) compared to WT littermates (**Fig. 5b-j**).

To further explore how these gene expression changes might have happened, we next performed transcription factor (TF) target enrichment analysis to identify potential upstream TFs responsible for observed changes in gene expression of Nutlin-3-treated *Fmr1* KO mice. Our analysis showed that the top TFs are mainly involved in stem cell fate specification (such as MEOX1, MEOX2, PRRX1, BNC2, SOX18 and TWIST1) [47–51] and/or extra cellular matrix organization (such as HEYL, TBX18, PRRX1, PRRX2 and TCF21)[52–54] (**Fig. 6a;; Additional file 3: Fig. S3**). We then assessed the relationship among our top TFs using a published TF network. We found that our top TFs related to Nutlin-3 treatment in KO mice showed a high level of interactions (**Fig. 6b**). Together, our transcriptomic analysis of the hippocampal tissue supports that the long-lasting rescue effects of Nutlin-3 treatment on impaired adult neurogenesis and dependent cognitive functions of *Fmr1* KO mice might be through modulating the adult neural stem cell niche.

**Fig. 6:**
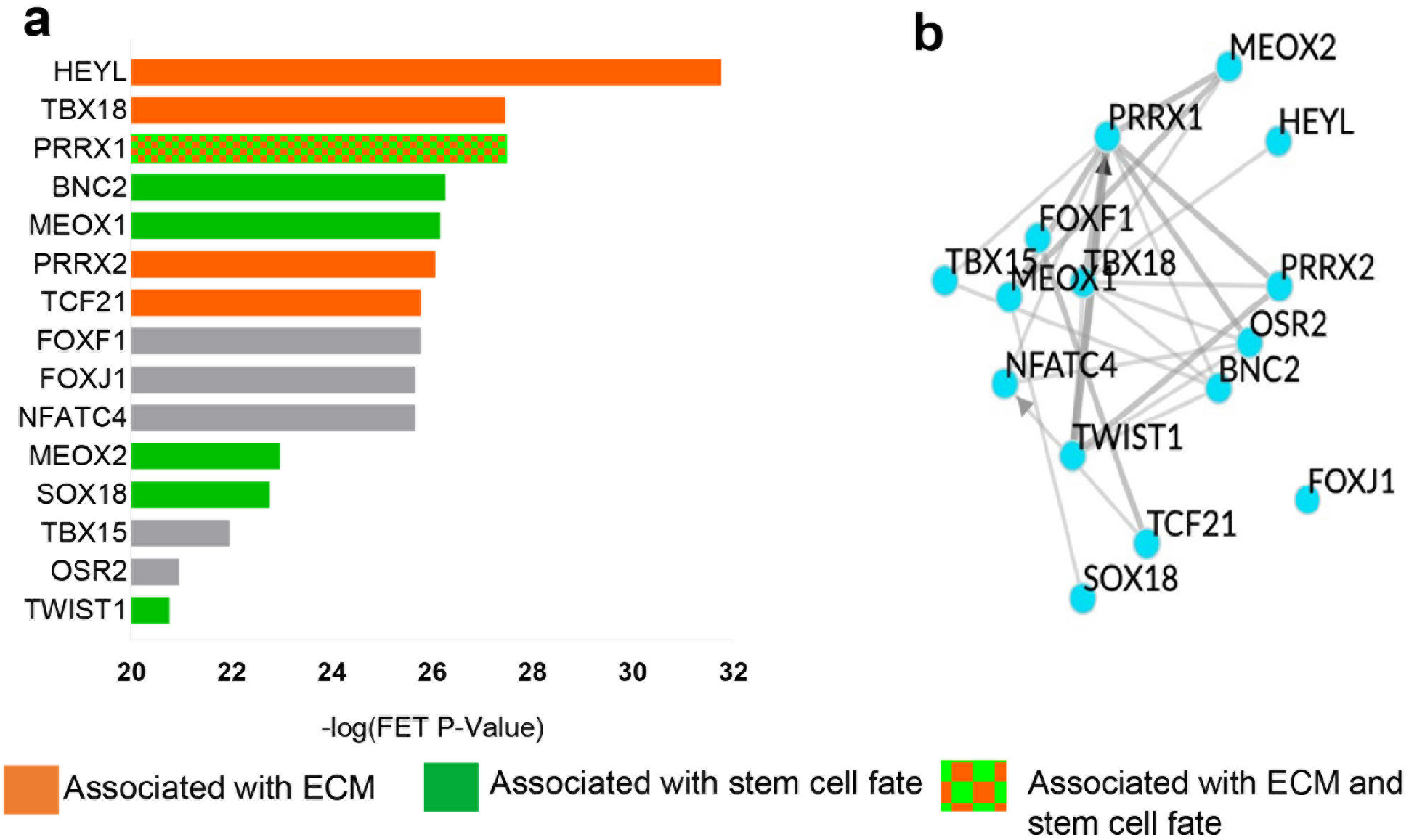
Top TFs ranked by TFs enrichment analysis were associated to ECM and stem cell fate. **a** Transcription factor target enrichment analysis of differentially expressed genes in *Fmr1* KO treated with Nutlin-3 vs vehicle using average integrated ranks across all libraries through ChEA3 (n = 3). FET, Fisher’s exact test. Orange bars indicate TFs associated with ECM. Green bars indicate TFs associated with stem cell fate. Orange and green patterned bars indicate TFs associated to both ECM and stem cell function. **b** top TFs network generated by Chea3 using average integrated ranks across all libraries.

## Discussion

This study sought to test the hypothesis that transient therapeutic intervention can produce long-lasting beneficial effects on cognitive functions in a mouse model of FXS. Our results demonstrate that a transient Nutlin-3 treatment of young adults for 10-days restored impaired hippocampal neurogenesis and related cognitive abilities in *Fmr1* KO mouse for at least 4 months after treatment cessation. Together with our publications [19, 20], these findings indicate not only that brief Nutlin-3 treatment rescues the neurogenic and cognitive deficits in adult FXS mice, but also that these beneficial effects are sustained long after the end of treatment. Our data also suggest that Nutlin-3 treatment during early adulthood time window might establish the normal adult NSC niche required for intact neurogenesis and cognitive performances in the absence of FMR1.

Numerous therapeutic alternatives including newly developed compounds or repurposed drugs have been proposed for FXS [19, 55]. There are many advantages of drug repurposing in the treatment of disease, including shortening the time frame and reduced cost associated with new drug development [56]. When assessing the feasibility of initiating treatments, an obvious concern is the resulting toxicity from long-term administration. For these reasons, there has been extensive interest in the possibility of repurposing drugs with potentially long-lasting therapeutic effects. However, only very few studies have assessed persistent effect of treatment long after treatments are stopped. In a recent study, minocycline treatment effect has lasted for 4 weeks in young FXS mice but not in adult FXS mice [14]. In another study, transient treatment of FXS rats with lovastatin at 4 weeks of age for 5 weeks prevented the emergence of cognitive deficits in object-place recognition and object-place-context recognition[11]. The authors show that corrective effect has sustained for at least 3 months (the last time point tested) after treatment termination and the observed restoration of normal cognitive function is associated with sustained rescue of both synaptic plasticity and altered protein synthesis[11]. One promising candidate for drug reproposing is a group of MDM2 inhibitors and its prototype is Nutlin-3. Nutlin-3 is a small molecule that specifically inhibits MDM2, an E3 ubiquitin ligase, and the best known MDM2 targets is tumor suppressor TP53, therefore Nutlin-3 and its derivative have been worked on extensively and used in clinical trial for cancer treatment [57]. Our lab has found that, in adult NSPCs, FMR1 directly regulates the expression levels and activities of MDM2, which targets TP53 and HDAC1 [19, 20]. Our published studies have shown that Nutlin-3 administration at a dosage significantly lower than those used for cancer treatment rescues impaired hippocampal neurogenesis and cognitive functions in either 2-month-old young adult FXS mice or 6-month-old mature adult FXS mice analyzed shortly after the treatment [19, 20]. However, the long-lasting effect of Nutlin-3 was unknown. Our current study has addressed this important question and taken one step further to potential therapeutic applications of MDM2 inhibition for treatment of FXS.

Understanding the molecular mechanism underlying drug action is important for both therapeutic application and improvement of drug development. To investigate the mechanism underlying the long-lasting effect of Nutlin-3, we first determined whether this effect was due to persistent changes in intrinsic properties of NSCs by using primary NSPCs isolated from *Fmr1* KO or WT mouse hippocampus. We have previously shown that NSPCs isolated from 2-month-old *Fmr1* KO hippocampus had reduced *TP53* gene expression, increased proliferation, and reduced neuronal differentiation, which can be corrected by Nutlin-3 treatment [19]. *TP53* gene encodes a transcription factor TP53 regulating a network of target genes that play roles in various cellular processes including but limited to apoptosis, cell cycle arrest, genomic integrity, metabolism, redox biology and stemness [58]. p53 binds DNA in a sequence-specific manner and recruits transcriptional machinery components to activate or suppress expression of a network of target genes [59]. TP53 has also been shown to regulate gene expression through epigenetic mechanisms [60–62], which may lead to long-lasting alteration in gene expression. We therefore hypothesized that Nutlin-3 treatment may exert sustained therapeutic effect on FXS mouse model through modulating epigenetic pathways in NSPCs. To our surprise, our results indicate that transient Nutlin-3 treatment did not lead to persist corrections in active MDM2 levels nor proliferation and differentiation of *Fmr1* KO NSPCs. This suggests that, unlike the immediate response to Nutlin-3 treatment, the long-lasting therapeutic effect of Nutlin-3 on neurogenesis might not be mediated through modulating NSPC intrinsic properties [19, 20]. Because adult neurogenesis is regulated by both NSC intrinsic pathways and extrinsic stem cell niche [63], we performed gene expression profile analysis of *Fmr1* KO and WT hippocampal tissue. Nutlin-3 treatment did not change the gene expression profile of WT hippocampus which is supported by our previous findings [19, 20]. On the other hand, Nultlin-3-treated KO mice mounted persistent and significant gene expression changes compared to vehicle-treated KO mice and WT mice. Among DEGs, we found mRNAs of proteins associated to extra cellular matrix, cell membrane and secreted factors, many of which have been shown to regulate adult neurogenesis [42–44]. For example, genes in TGFβ and BMP signaling are upregulated in Nutlin-3-treated KO mice, which we have confirmed using qPCR. It has been shown that TGFβ and BMP activation in adult NSC niche can activate adult neurogenesis [64, 65]. In addition, *Igf2* mRNA expression levels were significantly higher in Nutlin-3-treated KO hippocampus compared to either Vehicle-treated KO hippocampus or WT mice. IGF2 which has also been shown to promote adult NSC proliferation and neurogenesis[46].

One potential limitation of this study is that we have not defined whether an age range or a critical period exists for the initial Nutlin-3 treatment to achieve the long-lasting effectiveness of Nutlin-3 treatment. Future experiments on Nutlin-3 administration time at younger or older ages than 2 months should be considered. In addition, we showed that the beneficial effects of Nutlin-3 on impaired neurogenesis and cognition of FXS mice sustained for at least 4 months. Whether the effect lasts for a longer period or even for the rest of the animal’s life will need to be addressed in future studies. Furthermore, we have assessed adult neurogenesis-dependent behaviors. It is possible that Nutlin-3 also improves other aspects of behavioral deficits in FXS mice which is independent of adult neurogenesis, therefore it will be beneficial to assess whether the beneficial effects of Nutlin-3 can be generalized to other forms of cognitive and behavior functions found in FXS. Finally, although our transcriptomic analysis has provided important clue for the long-term effect of Nutlin-3 treatment, a comprehensive assessment of gene expression and epigenetic profiles of neurogenic niche will be needed to fully understand the molecular basis of persistent effect of Nutlin-3.

## Conclusions

In summary, our findings indicate that a brief Nutlin-3 treatment of young adult FXS mice has a long-lasting therapeutic effect on both neurogenesis and behaviors and that the sustained beneficial effect might be exerted through modulating adult NSC niche. Our observation strengthens the idea that Nutlin-3 is one of the ideal candidates for optimal therapy with the minimal toxicity in a targeted therapeutic approach. The findings provide proof-of-concept evidence that FXS, and perhaps neurodevelopmental disorders more generally, may be amenable to transient, early intervention to permanently restore normal cognitive functions.

## Abbreviations

ASD: autism spectrum disorder
ANOVA: Analysis of variance
BrdU: Bromodeoxyuridine
cKO: Conditional knockout
DEG: Differentially expressed gene
CE: Coefficient of error
DG: Dentate gyrus
ECM: Extracellular matrix
Erdr1: Erythroid differentiation regulator 1
Fig: Figure
*Fmr1*: Fragile x mental retardation protein 1
FMR1: Translational regulator 1 (official name)
FMRP: Fragile x mental retardation protein (previous name)
FXS: Fragile x syndrome
GAPDH: Glyceraldehyde-3-phosphate dehydrogenase
GC: Granule cell
GFP: Green fluorescent protein
GFAP: Glial fibrillary acidic protein
GO: Gene ontology
HDAC1: Histone deacetylase 1
KO: Knockout
qPCR: quantitative polymerase chain reaction
MCM2: Minichromosome maintenance complex component 2
MDM2: mouse double minute 2
NeuN: Neuronal nuclei antigen
NIH: National institute of health
NLR: Novel location recognition
NOR: Novel object recognition
NSPC: Neural stem/progenitor cell
NSC: Neural stem cell
NP: Neural progenitors
Nut3: Nutlin3
p53: Protein
PFA: Paraformaldehyde
RBFOX3: RNA binding fox-1 homolog 3
RGL: Radial glial like
SGZ: Subgranular zone
SVZ: Subventricular zone
TF: Transcription factor
Veh: Vehicle
WT: Wild type

## ACKNOWLEDGMENTS

We thank Y. Xing, K. Schoeller, J. Le, and ? for technical assistance, J. Panksepp, D. Bolling and MM Eastwood, K. Knobel at the Waisman IDD Model Core, UW-Madison Biotechnology Center for next generation sequencing services. This work was supported by grants from the National Institutes of Health (R01MH118827, R01MH116582, R01NS105200 to X.Z., R01NS064025, R01AG067025, U01MH116492 to D. W., U54HD090256 and P50HD105353 to the Waisman Center), Jenni and Kyle Professorship to XZ, Wisconsin Distinguished Graduate Fellowship to SJ

## AUTHOR CONTRIBUTIONS

XZ conceived the concept. XZ, SJ. YL designed and performed experiments, collected data, and analyzed data. SJ and XZ wrote the manuscript. JS and DW performed bioinformatics analysis

## Supplementary File List

**Additional file 1:**

**Code S1**. R codes for differentially expressed gene analysis.

**Code S2**. R codes for STAR alignment

**Additional file 2:**

**Table S1**. Primer sequences for qPCR.

**Table S2**. Result of RNA-seq read alignment.

**Table S3**. Raw read counts of RNA-seq samples

**Table S4**. Differentially expressed genes among experimental groups.

**Table S5**. GO enrichment analysis results

**Additional file 3:**

**Fig. S1**. Transient treatment with Nutlin-3 has long-lasting rescue effect on cognitive deficits in mice with selective deletion of Fmr1 in adult new neurons

**Fig. S2**. Boxplot to show the density distribution of raw log-intensities of RNA-seq data of all samples.

**Fig. S3**. GO analysis of top upstream TF.

