## supplementary Figures for "Sustained correction of hippocampal neurogenic and cognitive deficits after a brief treatment by Nutlin-3 in a mouse model of Fragile X Syndrome"

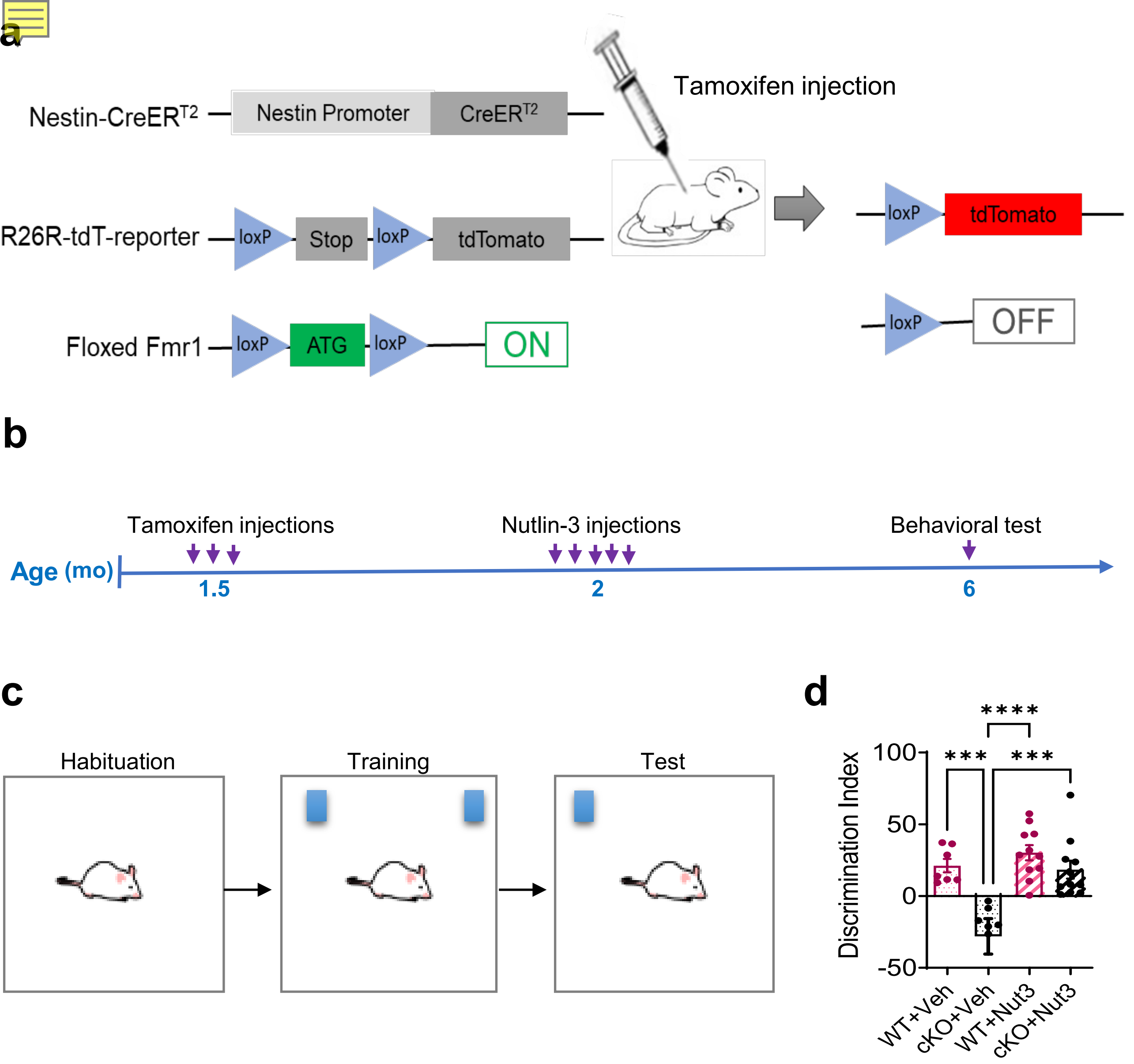


**FigureS1: Transient treatment with Nutlin-3 has long-lasting rescue effect on cognitive deficits in mice with selective deletion of Fmr1 in adult new neurons. a** An inducible FMRP conditional knockout mouse line was created by crossing Nestin-CreER$^{T2}$ mice, ROSA26-STOP-tdTomato (Ai14) mice and *Fmr1* floxed (*Fmr1* cKO) mice. Administration of tamoxifen to adult mice results in the removal of the first exon of the mouse *Fmr1* gene and the "Stop" codon before tdTomato (tdT) in Nestin-expressing cells and their subsequent progenies. **b** Experimental scheme for analyzing cognitive performances in *Fmr* cKO treated with Nutlin-3 or vehicle **c** Schematic of novel location recognition test for assessing spatial memory in *Fmr1* cKO mice. **d** Nutlin-3 treatment fully rescued spatial memory deficits in *Fmr1* cKO mice 4 months after injection (n = 6 to 11 mice per group). ***P < 0.001. Data are presented as means ± SEM.

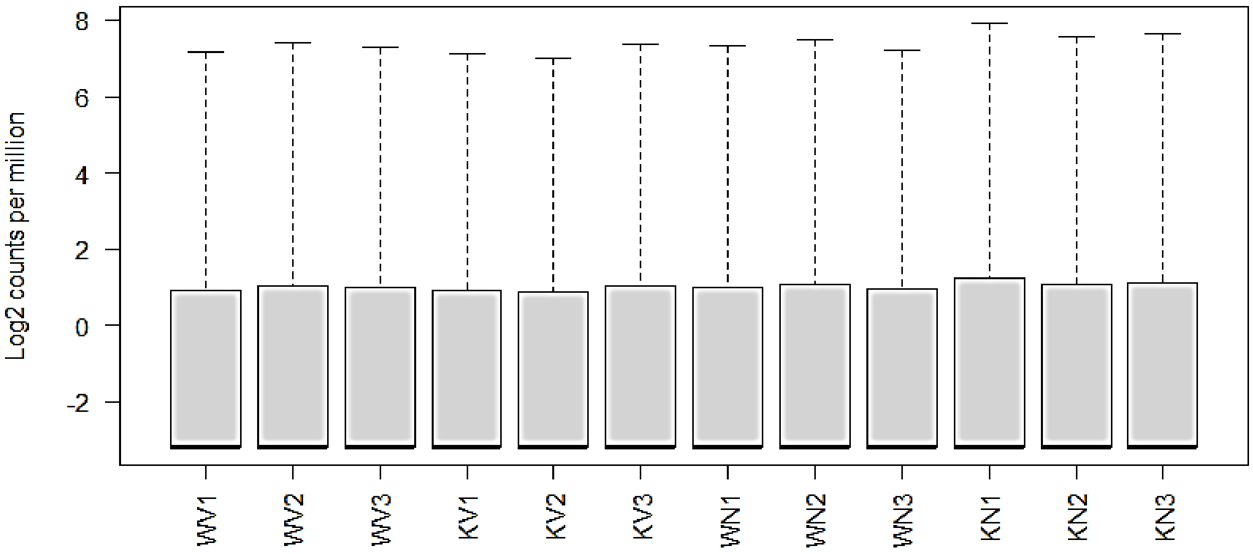


**Figure S2: Boxplot to show the density distribution of raw log-intensities of RNA-seq data of all samples.**

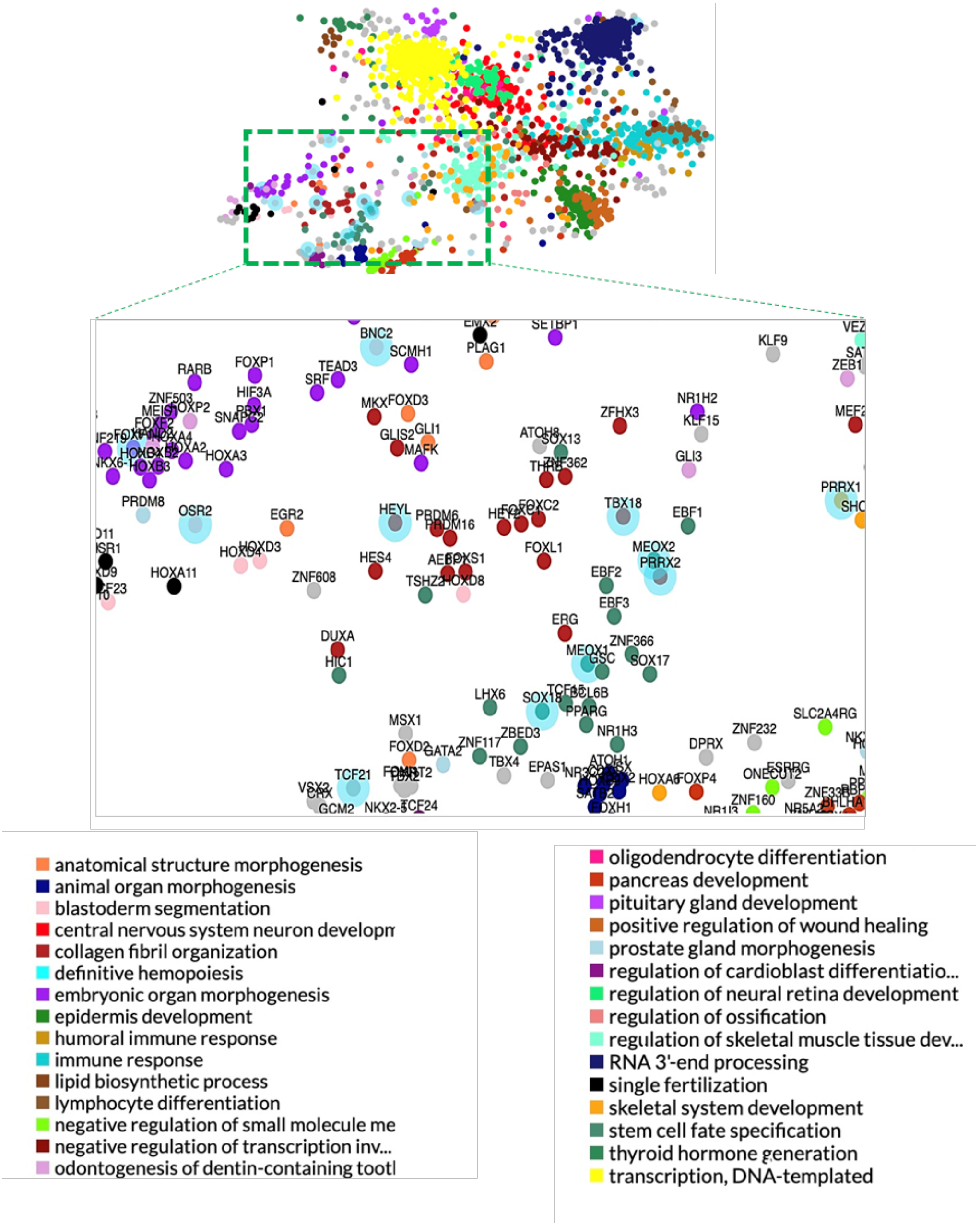


**Figure S3**. **GO analysis of top upstream TF.**

ChEA3-generated GO enrichment analysis of top upstream TFs ranked by ChEA3, using average integrated ranks across all libraries, for DEGs between Fmr1 KO treated with Nutlin-3 vs vehicle.
